# PrimerServer: a high-throughput primer design and specificity-checking platform

**DOI:** 10.1101/181941

**Authors:** Tao Zhu, Chengzhen Liang, Zhigang Meng, Yanyan Li, Yayu Wu, Sandui Guo, Rui Zhang

## Abstract

**Summary:** Designing specific primers for multiple sites across the whole genome is still challenging, especially in species with complex genomes. Here we present PrimerServer, a high-throughput primer design and specificity-checking platform with both web and command-line interfaces. This platform efficiently integrates site selection, primer design, specificity checking and data presentation. In our case study, PrimerServer achieved high accuracy and a fast running speed for a large number of sites, suggesting its potential for molecular biology applications such as molecular breeding or medical testing.

**Availability and Implementation:** Source code for PrimerServer is available at https://github.com/billzt/PrimerServer. A demo server is freely accessible at https://primerserver.org, with all major browsers supported.

## 1 Introduction

Designing specific primers for templates such as genomic DNA or cDNA is an important step in PCR. The genomes of many species, especially angiosperms, contain large amounts of repetitive elements, expanded gene families, and, in polyploids, homologous sub-genomes. This high degree of sequence similarity within a genome can result in primer binding to unintended regions, leading to false-positive amplicons. Thus, it is necessary to integrate a specificity checking step into primer designing tasks. Although several tools, such as Primer-BLAST (Ye *et al.*, 2012), PolyMarker (Ramirez-Gonzalez *et al.*, 2015), GSP (Wang *et al.*, 2016), and ThermoAlign (Francis *et al.*, 2017), have been developed to design specific primers, they have somewhat less power when dealing with high-throughput jobs. Here, we present PrimerServer, an efficient platform with both a web and command line interface, to design specific primers for multiple sites simultaneously.

## 2 Methods

### 2.1 Workflow

The workflow of PrimerServer consists of three steps (Figure 1): primer design, primer specificity check, and data integration. In the first step, PrimerServer accepts user-inputted target site information, uses Samtools (Li *et al.*, 2009) to extract corresponding template fragments and then passes these fragments to Primer3 (Untergasser *et al.*, 2012) to obtain a list of candidate primer pairs. In the second step, the sequences of these candidate primer pairs are extracted and used as queries in BLAST (Camacho *et al.*, 2009) searches against background sequences specified by the user (e.g., whole genomes, transcriptomes, or other specific sets of sequences). The high-scoring segment pairs (HSPs) obtained from BLAST are paired and filtered by their coordinates to generate possible PCR amplicons within a size range specified by the user. Next, the melting temperatures (Tm) of these paired HSPs are calculated and filtered using the method of Francis *et al.* (2017). Finally, for each target site, PrimerServer first sorts the potential primer pairs by the number of possible PCR amplicons and then by the penalty score from Primer3, and displays them in a pretty-formatted web page. In addition, during the second step, PrimerServer can also accept user-provided primers and output possible amplicons without running the whole pipeline.

**Figure 1.**
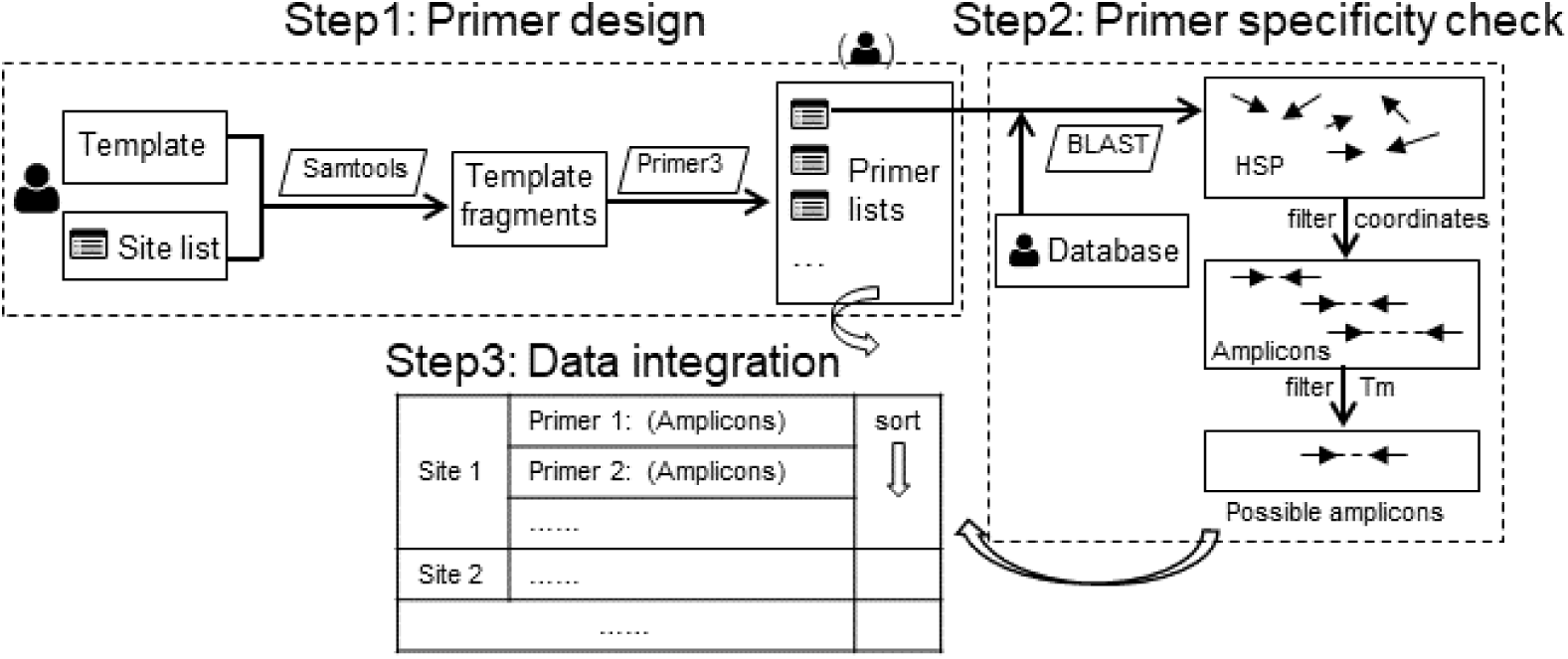
Workflow of PrimerServer. Rectangles indicate input and output files. Rhomboids indicate external programs.

### 2.2 Web implementation and user interface

PrimerServer provides a user-friendly interface for selecting databases, inputting target sites, setting parameters, and viewing or saving results. It contains three modules, and the user can easily switch between modules by clicking on the navigation tabs (Supplemental Fig. S1-3). The first module “Design & Check” accepts a list of user-inputted sites, runs the whole pipeline, and returns the appropriate primer pairs with amplicon numbers marked. For each site, PrimerServer draws an interactive graph indicating the positions of the target region and its primers using the D3.js library. The second module “Specificity Check” directly accepts user-provided primers and returns possible amplicons for each primer group. In both modules, users can download the results as text files to their local devices, or they can save the graphic HTML results in the webserver, which are listed in the third module “Recently Saved Results”.

The web interface was developed using the Twitter Bootstrap framework based on HTML5 and JavaScript. User-specified parameters are remembered and stored in their browsers and can automatically be applied at the time of the next visit. Our demo server is hosted on a SuperMicro® workstation equipped with 80 Intel® Xeon® CPUs (2.40GHz), 128 GB RAM, and running the CentOS 6.9 system. Currently 18 species are included in the demo server. New PrimerServer instances can be easily deployed on any Linux computer with web service and a PHP environment.

## 3 Results

As a case study, we used PrimerServer to design specific primers for genome-wide insertion-deletion (INDEL) polymorphic sites in the allotetraploid cotton (*Gossypium*), whose genome consists of ≥60% repetitive elements (Zhang *et al.*, 2015). Among the 14,754 INDEL sites (≥20bp) inferred from a comparison between *G. hirsutum* and *G. barbadense*, we successfully designed unique primers for 1,559 sites. PCR validation of 50 randomly selected primers revealed that 84% are unambiguously genome-wide unique (Supplemental Text).

## 4 Conclusion

PrimerServer is an efficient platform for designing high-throughput specific primers. According to our case study, it performs exceptionally well, even in polyploids with a large number of repeat sequences. We anticipate that PrimerServer will be useful in molecular breeding, medical testing, and many other fields.

